# Nanocondensate tunnelling

**DOI:** 10.64898/2026.06.22.731667

**Authors:** Luyi Han, Yue Wan, Zhenxi Guo, Shuo Gao, Ningli Wang, Yingyan Mao

## Abstract

Nanocondensates occupy mesoscopic scale in both spatial scale and thermodynamic state, and the mesoscopic scale is central to structure-function conversion in living systems. However, how the dual mesoscopic properties of nanocondensates support biological function remains unclear because their small size and rapid dynamics hinder direct observation in living cells. Here we construct nanocondensates in situ at the cell-membrane interface and observe them permeating membranes within seconds, diffusing freely, exiting cells within minutes, re-entering neighbouring cells and traversing multilayered barriers. We report the first observation of this phenomenon and term it nanocondensate tunnelling. We find that nanocondensates cross membranes without membrane enclosure, preserving their wetting characteristics after passage. Theoretical and ultrastructural analyses support a transient-pore model linking this behaviour. We exploit nanocondensate tunnelling to transport small molecules, siRNA and antibodies across cellular barriers, and small molecules and siRNA across a multilayered corneal barrier, demonstrating its functional capacity for intercellular transfer. Our results suggest that nanocondensate tunnelling represents a previously undescribed mode of intercellular transfer. Nanocondensate tunnelling provides a physical basis for biologics delivery and a testable framework for investigating unresolved biological phenomena such as viral permeation.

## Introduction

The mesoscopic scale is central to structure-function conversion in living systems^1,2^. From a renormalization-group (RG) perspective^3,4^, it is an effective scale at which microscopic perturbations have not yet been completely averaged out and collective order begins to emerge. In living matter, this scale is defined not only by spatial scale but also by the thermodynamic state of molecular assemblies^5^. At mesoscopic spatial scales, local conformational changes can reshape the overall response of an interaction domain while preserving component organization^6,7^, as in chaperone-mediated remodeling of client proteins^8^ and gene regulation by transcriptional complexes^9^. Along the thermodynamic dimension, biomolecular condensation can generate mesoscopic states in which reversible molecular interactions and continuous component exchange maintain fluidity while preserving collective organization^10^, as exemplified by synaptic-vesicle clustering and the layered organization of nucleolar subcompartments^11,12^. Coupling between spatial and thermodynamic mesoscopic properties can give rise to assemblies such as nanocondensates, whose dynamic yet persistent material state permits rapid component exchange and molecular information transfer while sustaining functional output, suggesting important biological roles^13^. However, their small size and rapid dynamics make nanocondensates difficult to observe and characterize directly in living cells, limiting understanding of how their mesoscopic physical properties support biological function^14^.

Here we construct nanocondensates in situ at the cell-membrane interface through sequential cell-penetrating peptide (CPP) adsorption and CPP-silk fibroin (SF) co-condensation and observe them permeating the plasma membrane within seconds. Once in the cytoplasm, they exhibit diffusion-like motion and remain free of a detectable persistent membrane enclosure. They exit individual cells within minutes and subsequently enter neighbouring cells. Through successive cycles of membrane entry and exit, these nanocondensates traverse multilayered tissue barriers sealed by tight junctions. We report the first discovery of this phenomenon and term it **nanocondensate tunnelling**. Nanocondensate tunneling differs from the prevailing understanding of cellular transmembrane transport: nanoscale dimensions and the condensate phase state jointly drive rapid transmembrane permeation without membrane enclosure^15^, allowing the nanocondensates to retain their wetting characteristics^15,16^ after permeation and thereby enabling transmembrane behavior to proceed as a continuous process. Computational analyses support a transient-pore model linking this behaviour to nanoscale dimensions and a fluid condensate state. Our results suggest that nanocondensate tunnelling represents a previously undescribed mode of intercellular material transfer. It provides a physical basis for biologics delivery and a testable framework for investigating unresolved biological phenomena such as viral permeation^17,18^.

### Spatial segregation of cohesion and surface stabilization defines the nanocondensate state

To establish a physically defined nanocondensate state for cellular analysis, we first asked what prevents condensates from dissolving, coarsening into micrometre-sized droplets or solidifying^19,20,21^. We modelled fusion between equal droplets with radii ranging from 50 nm to 2 µm (Fig. 1a). As radius increased, the Brownian encounter frequency decreased as R⁻³, whereas the magnitude of capillary free-energy release upon fusion increased as R²^22,23^ (Fig. 1b,c). Surface charge introduced an opposing approach barrier proportional to Rψ², where ψ denotes the surface potential^24^. In the model, surface potentials of 10-25 mV consequently suppressed successful fusion (Fig. 1d,e). Stable nanocondensates therefore required sufficient cohesion to resist dissolution and sufficient surface repulsion to suppress coarsening^25,26^. A dimensionless state map combining non-electrostatic cohesion, charge-dependent repulsion and network solidification identified a restricted nanocondensate window between dispersed, coarsening and gelled states (Fig. 1f). This window narrowed as the target diameter decreased (Fig. 1g), predicting that internal cohesion and surface stabilization must be spatially separated and balanced with increasing precision at smaller sizes. Detailed calculations and descriptions are provided in Supplementary Data1.

**Fig. 1.**
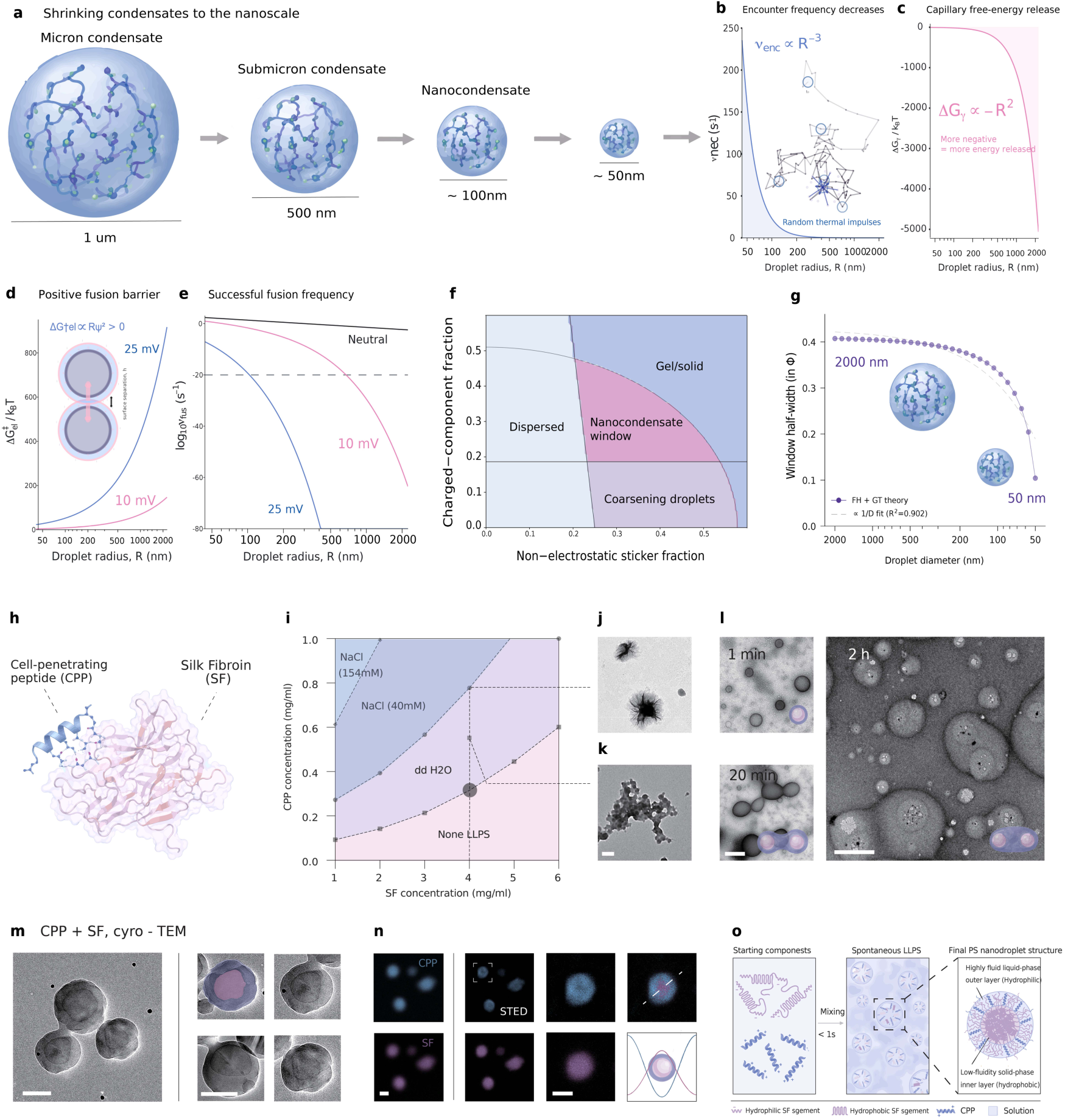
Spatial segregation of cohesion and surface stabilization defines the nanocondensate state. a, Schematic of condensate miniaturization. **b,c,** Predicted dependence of Brownian encounter frequency (b) and capillary free-energy change upon fusion (c) on droplet radius, R. **d,e,** Predicted electrostatic approach barrier (d) and successful fusion frequency (e) for neutral droplets or droplets with surface potentials of 10 or 25 mV. **f,** Dimensionless state map showing dispersed, coarsening, nanocondensate and gelled or solid states. **g,** Nanocondensate-window half-width, Φ, versus droplet diameter. Points show model predictions; the dashed line shows a reciprocal-diameter fit (R² = 0.902). **h,** Structural representations of cell-penetrating peptide (CPP) and silk fibroin (SF). **i,** CPP-SF phase diagram at the indicated NaCl concentrations. Binary-component stoichiometry (BcS) denotes the CPP-to-SF mass ratio. **j,k,** TEM images of fibrillar (j) and irregular aggregated (k) assemblies at higher BcS. **l,** Time-course TEM images showing nanocondensate fusion. **m,** Cryo-TEM images showing a core-shell architecture. **n,** Conventional and STED microscopy showing peripheral CPP enrichment and an SF signal extending across the condensate and peaking towards its centre; right, radial intensity profiles. **o,** Proposed model in which a cohesive core enriched in hydrophobic SF segments is surrounded by a hydrated shell containing hydrophilic SF segments and associated CPP. Phase boundaries were reproduced in three independent preparation series. Images are representative of three independently prepared samples. No null-hypothesis test was applied to theoretical panels. Scale bars, 100 nm (j-n).

Guided by this design principle, we combined a positively charged, conformationally constrained cell-penetrating peptide (CPP)^27^ with amphiphilic silk fibroin (SF)^28,29^ (Fig. 1h). We defined the mass-based binary-component stoichiometry (BcS) as the CPP-to-SF mass ratio. CPP and SF co-condensed only within a narrow BcS range, which shifted towards higher CPP concentrations as salinity increased (Fig. 1i). At BcS = 1:15, turbidity reached an OD₆₀₀ of approximately 0.5, but centrifugation recovered only about 5% of the material. The resulting assemblies had diameters concentrated between 70 and 100 nm and contained moderate levels of SF β-sheet structure (Extended Data Fig. 1a-d). Increasing BcS raised the centrifugally recovered fraction to approximately 22% at 1:12 and 38-40% at 1:8. Particle size and SF β-sheet content increased concurrently. Low-to-intermediate-BcS assemblies were spherical and underwent fusion, whereas higher BcS conditions produced irregular aggregates and fibrils (Fig. 1j-l). Desolvation collapsed the hydrated assemblies into smaller nanospheres, and FRET confirmed molecular proximity between CPP and SF (Extended Data Fig. 1e,f). Together, these measurements resolved a BcS-dependent progression from fluid nanocondensates to increasingly solid-like assemblies and fibrils (Extended Data Fig. 1g).

We next tested how elevated ionic strength altered the CPP-SF assembly window. At 154 mM NaCl, the phase boundary shifted to peptide concentrations approximately tenfold higher than those required in water. Tandem duplication of CPP (dCPP) nevertheless supported co-condensation under this condition, indicating that increased local multivalency partly compensated for electrostatic screening (Extended Data Fig. 2a). By contrast, 10 mM FeCl₃ or CaCl₂ strongly restricted phase separation, and Fe³⁺ disrupted the ordered CPP conformation^30,31^ (Extended Data Fig. 2b,c). Together, these perturbations showed that electrostatic recruitment depended on both local charge density and the conformational organization of charged residues.

We next examined how CPP and SF were spatially organized within the nanocondensates. Cryo-TEM revealed comparable core-shell architectures in CPP-SF and dCPP-SF nanocondensates (Fig. 1m and Extended Data Fig. 2g). Upon desolvation, the apparent shell thickness decreased from approximately 35 nm to less than 10 nm, whereas the 50-60 nm core remained largely unchanged (Extended Data Fig. 2d). Surface-weighted Raman spectroscopy detected protonated amino groups, and the condensates exhibited slightly positive ensemble ζ-potentials of approximately +2 to +5 mV (Extended Data Fig. 2e,f). STED microscopy showed peripheral enrichment of CPP, whereas the SF signal extended across the condensate and peaked towards its center in both CPP-SF and dCPP-SF nanocondensates (Fig. 1n and Extended Data Fig. 2h). These orthogonal measurements revealed a hydrated shell with peripheral CPP enrichment surrounding an SF-containing condensate interior.

Together, the structural and perturbation data supported a model in which cationic CPP associates with anionic groups on SF, partially compensating the negative charge of SF and reducing electrostatic repulsion between SF molecules. In this model, local SF enrichment strengthens hydrophobic association and favours β-sheet formation, generating a cohesive core enriched in hydrophobic SF segments. Hydrophilic SF segments and associated CPP form a hydrated outer shell. Residual positive charge on shell-localized CPP is consistent with the slightly positive ensemble ζ-potential and, together with shell hydration, would oppose further coalescence. This spatial separation of core cohesion from shell-mediated electrostatic and hydration stabilization provides a mechanistic basis for the narrow BcS window that preserves nanoscale dimensions and fluidity (Fig. 1o). These results established a physically defined, BcS-tunable nanocondensate state for subsequent analysis in living cells.

### BcS-defined membrane-confined nanocondensates undergo rapid cellular entry

Having defined the BcS-dependent nanocondensate state in bulk, we next sought to reconstruct this state at a living membrane interface and examine its cellular behaviour^32^. CPP served as the membrane-localizing component because it contributed to co-condensation and exhibited intrinsic membrane affinity^33^. Brief exposure at low micromolar concentrations produced a broadly distributed CPP signal along the plasma membrane. CPP alone produced no detectable particulate internalization, whereas SF alone generated little intracellular particulate signal. By contrast, adding SF to CPP-decorated membranes at a nominal BcS of 1:14 reorganized membrane-associated CPP into puncta that colocalized strongly with SF (Fig. 2a,b and Extended Data Fig. 3a,b). These observations supported in situ CPP-SF co-assembly at the plasma membrane. During this reconstruction, we unexpectedly observed SF-containing puncta inside the cells (Fig. 2a).

**Fig. 2.**
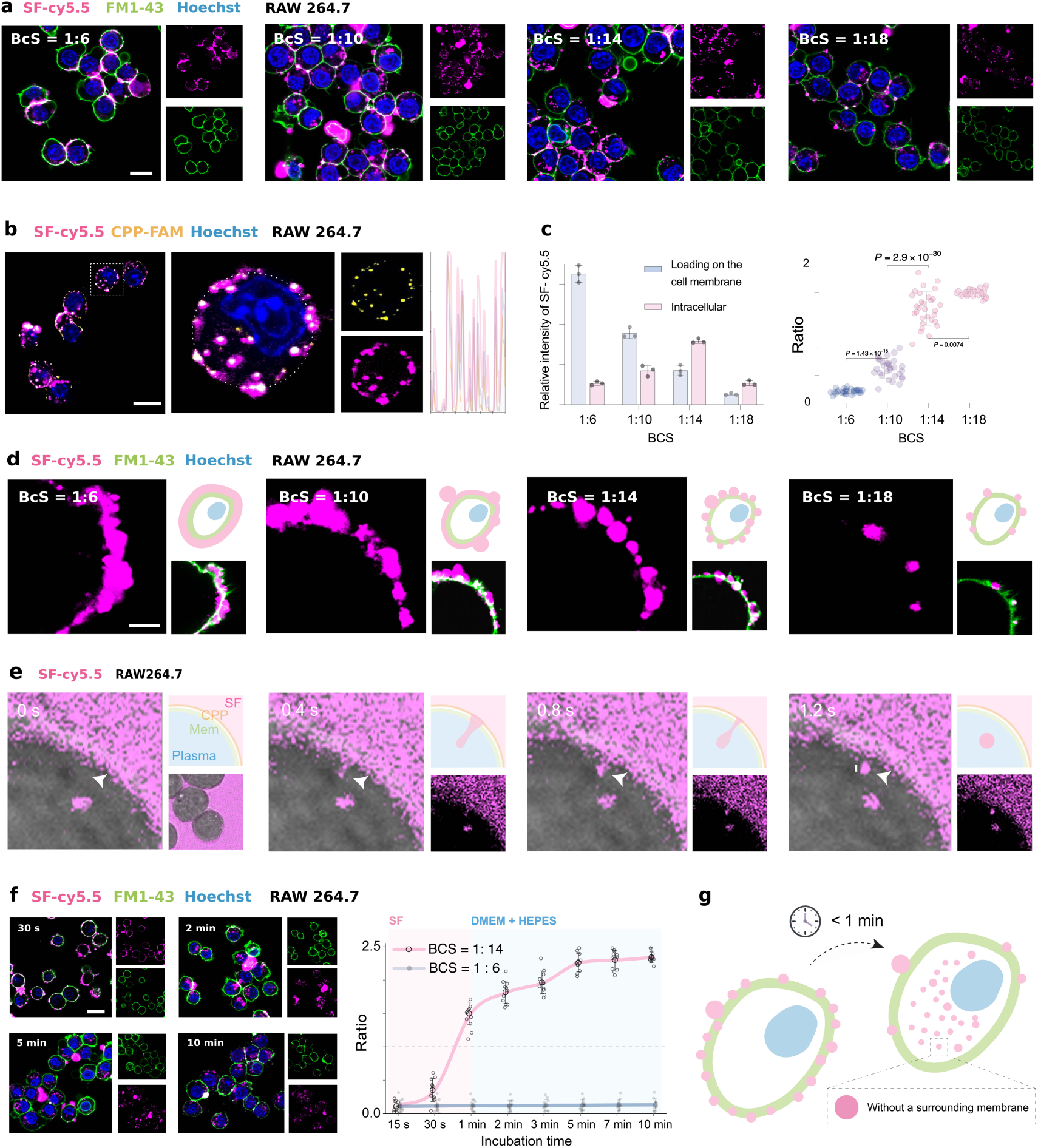
BcS-defined membrane-confined nanocondensates undergo rapid cellular entry. a, Confocal images of RAW 264.7 cells after 30 s of CPP loading and 1 min of SF incubation at the indicated BcS values. **b,** Colocalization of CPP-FAM and SF-Cy5.5 at BcS = 1:14. **c,** Quantification of membrane-associated and intracellular SF signals (left) and intracellular-to-membrane signal ratios (right). d, STED images of membrane-associated SF assemblies formed at 4 °C. **e,** Live-cell image sequence showing the inward translocation of a Cy5.5-labelled nanocondensate across the plasma membrane of a CPP-pretreated RAW 264.7 cell. Arrowheads mark the translocating nanocondensate. **f,** Representative confocal images and time-dependent intracellular accumulation of SF at BcS = 1:14 and 1:6. **g,** Schematic summarizing rapid nanocondensate entry. For c and f, n = 4 independent experiments; measurements from 30 cells per condition were averaged within each experiment. Data are mean ± s.e.m. of the experiment means. In c, groups were compared using two-sided one-way ANOVA followed by Holm-Šidák correction. Adjusted *P* values for BcS = 1:14 versus 1:6, 1:10 and 1:18 were 0.0008, 0.012 and 0.0016, respectively. In f, the time courses were analysed using a two-sided mixed-effects model with Geisser-Greenhouse and Holm-Šidák corrections; the time-by-BcS interaction gave *P* < 0.0001. Scale bars, 10 µm (a, b, f), 1 µm (d), 100 nm (e) and 10 µm (e, lower inset).

To determine whether this intracellular accumulation reflected CPP abundance or the assembled material state, we held the SF concentration constant and varied CPP loading. If CPP alone drove internalization, intracellular SF would be expected to increase monotonically with CPP abundance. Instead, a nominal BcS of 1:14 produced the highest absolute intracellular SF signal. The CPP-rich BcS of 1:6 generated high membrane loading but little intracellular signal (Fig. 2a,c). At the CPP-poor BcS of 1:18, both membrane-associated and intracellular signals were weak, despite a high intracellular-to-membrane signal ratio. Absolute particulate internalization therefore peaked within a restricted BcS range rather than increasing with CPP loading.

We next examined the membrane-associated assemblies before internalization. Cooling cells to 4 °C slowed membrane dynamics and retained the assemblies at the cell surface long enough for imaging^34,35^. STED microscopy showed that a BcS of 1:14 produced the most distinct discrete puncta (Fig. 2d). CPP-rich conditions produced more continuous or fibre-like surface assemblies, whereas the CPP-poor BcS of 1:18 yielded sparse puncta. After the cells were returned to room temperature, fluorescence recovery reached 57.3% at a BcS of 1:14. The corresponding recoveries were 5.6%, 31.6% and 2.1% at BcS values of 1:6, 1:10 and 1:18, respectively (Extended Data Fig. 3c). Thus, the greatest absolute intracellular accumulation coincided with discrete and dynamically recovering membrane-associated assemblies.

We then resolved the kinetics of this behaviour. At a nominal BcS of 1:14, intracellular SF accumulation increased sharply within 1 min and was largely complete by 5 min. By contrast, little accumulation occurred at a BcS of 1:6 (Fig. 2f). Across four independent experiments, 47 inward translocation events were detected in 31 cells. In a representative event, an extracellular punctum extended inwards and separated from the plasma membrane within 1.2 s (Fig. 2e and Supplementary Video 1). No detectable large-scale alteration of membrane morphology accompanied this event. Intracellular SF puncta also lacked detectable colocalized membrane-dye signal at the resolution of fluorescence imaging (Fig. 2a,f).

The relationship between surface assembly and intracellular accumulation extended across cell types. At the same nominal BcS of 1:14, intracellular SF accumulation varied among HCEC, RPE, SRA and BV2 cells alongside differences in surface CPP loading (Extended Data Fig. 3d-f). HCEC cells showed higher CPP adsorption, whereas BV2 cells showed lower adsorption. In HCEC cells, decreasing the CPP-to-SF feed ratio from 1:14 to approximately 1:28 increased intracellular SF accumulation, whereas a further decrease to 1:36 reduced it (Extended Data Fig. 3g,h). Together, these measurements showed that BcS-defined, fluid nanocondensates assembled at the plasma membrane and underwent rapid cellular entry (Fig. 2g).

### Rapid cellular exit and intercellular transfer complete nanocondensate tunnelling

The rapid cellular entry observed prompted us to continue imaging the nanocondensates and characterize their subsequent cellular behaviour. We tracked 30 cells in each of four independent experiments from 15 to 40 min after assembly. Intracellular SF fluorescence peaked at 15 min and declined thereafter. By contrast, the plasma membrane-associated signal increased, peaked at 30 min and subsequently decreased (Fig. 3a and Extended Data Fig. 3i). By 40 min, the intracellular signal was nearly undetectable, and the intracellular-to-membrane fluorescence ratio had declined continuously from 15 min onwards. CPP and SF remained highly colocalized within the detectable puncta throughout this period. These observations established a temporally ordered redistribution from an intracellularly enriched state towards the plasma membrane, while the two components remained associated at fluorescence resolution (Fig. 3a,b). This temporally ordered redistribution suggested that intact nanocondensates could return from the cytosol to the plasma membrane and subsequently move into the extracellular space.

**Fig. 3.**
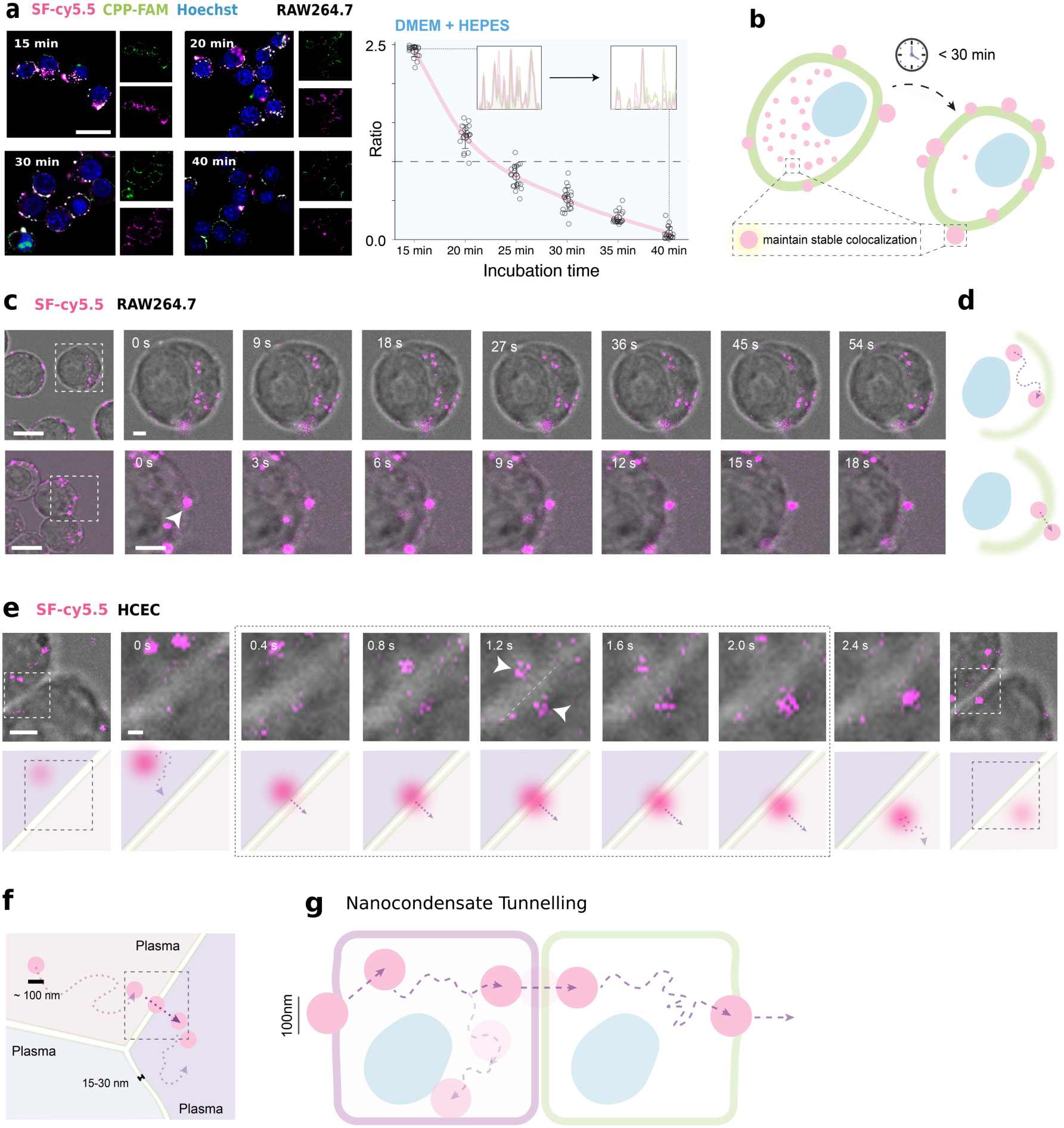
Rapid cellular exit and intercellular transfer complete nanocondensate tunnelling. a, Confocal images showing the time-dependent redistribution of CPP-SF nanocondensates in RAW 264.7 cells after cellular entry, together with the intracellular-to-plasma-membrane SF-Cy5.5 fluorescence ratio. Insets in the graph show representative CPP-FAM and SF-Cy5.5 intensity profiles. **b,** Schematic of intracellular nanocondensate redistribution towards the plasma membrane while CPP and SF remain colocalized. **c,** Live-cell sequence showing diffusion-like intracellular motion and outward translocation of an SF-Cy5.5-labelled nanocondensate in a RAW 264.7 cell. **d,** Schematic of nanocondensate exit from a cell. **e,** Live-cell sequence showing transfer of an SF-Cy5.5-labelled nanocondensate between adjacent HCEC cells; arrowheads indicate the transferring nanocondensate, and the diagrams below depict its position relative to the two cell boundaries. **f,** Geometrical model of a nanocondensate passing across two closely apposed plasma membranes. **g,** Integrated model of nanocondensate tunnelling, comprising cellular entry, intracellular motion, cellular exit and entry into a neighbouring cell. Images and quantification in a are from four independent experiments, with 30 cells analysed per time point in each experiment; cell-level measurements were averaged within each experiment. Data are mean ± s.e.m. Scale bars, 20 µm (a), 10 µm (c,e, overview images) and 1 µm (c,e time-lapse sequences).

We next characterized the intracellular puncta during this interval. After cellular entry, however, the puncta showed minimal colocalization with LysoTracker (Extended Data Fig. 5c). Together with the persistent CPP-SF colocalization, this observation showed that most detectable nanocondensates did not accumulate in acidic lysosomes during the observation period. Live-cell tracking at 10 min further revealed non-directed intracellular trajectories without persistent directional movement (Fig. 3c and Extended Data Fig. 5d). These trajectories were consistent with diffusion-like motion rather than sustained directional transport. During continued live-cell imaging, we captured nanocondensates moving outwards across the cell boundary. Across four independent experiments, 22 outward-translocation events were detected in 18 cells. In a representative event, an intracellular nanocondensate approached the plasma membrane, moved to its extracellular side over an 18-s sequence and remained transiently at the cell surface (Fig. 3c,d).

The observation of cellular exit raised the possibility that released nanocondensates could transfer directly into neighbouring cells. In confluent HCEC monolayers, adjacent plasma membranes can approach within approximately 15-30 nm, providing closely apposed cellular interfaces (Fig. 3e). Live-cell imaging captured 12 complete transfers across eight cell pairs in four independent experiments. In a representative event, a nanocondensate approached the intercellular interface from within one cell (Fig. 3f). Its fluorescence then extended across both sides of the interface and became approximately symmetric midway through the transfer. The signal subsequently resolved within the neighbouring cell, where the nanocondensate resumed diffusion-like movement. Redistribution across the interface occurred within approximately 1 s, with no resolvable pause between outward and inward translocation at the acquisition rate used. Together with the rapid entry observed, these observations revealed an integrated sequence of inward translocation, intracellular motion, cellular exit and neighbouring-cell entry. We define this behaviour as nanocondensate tunnelling (Fig. 3g). This raised a central mechanistic question: how can the same nanocondensate state repeatedly cross successive cellular interfaces?

### Mesoscopic properties support a transient-pore model of nanocondensate tunnelling

We next investigated the physical mechanism underlying nanocondensate tunnelling by analyzing membrane deformation at the condensate-membrane interface. Geometrical analysis revealed a strong dependence on particle size. At comparable wetting strength, smaller particles generated greater central curvature while distributing the deformation over fewer lipids. Reducing particle diameter from 160 to 80 nm doubled the total central curvature (Fig. 4a,b), concentrating leaflet-area mismatch and local membrane stress.

**Fig. 4.**
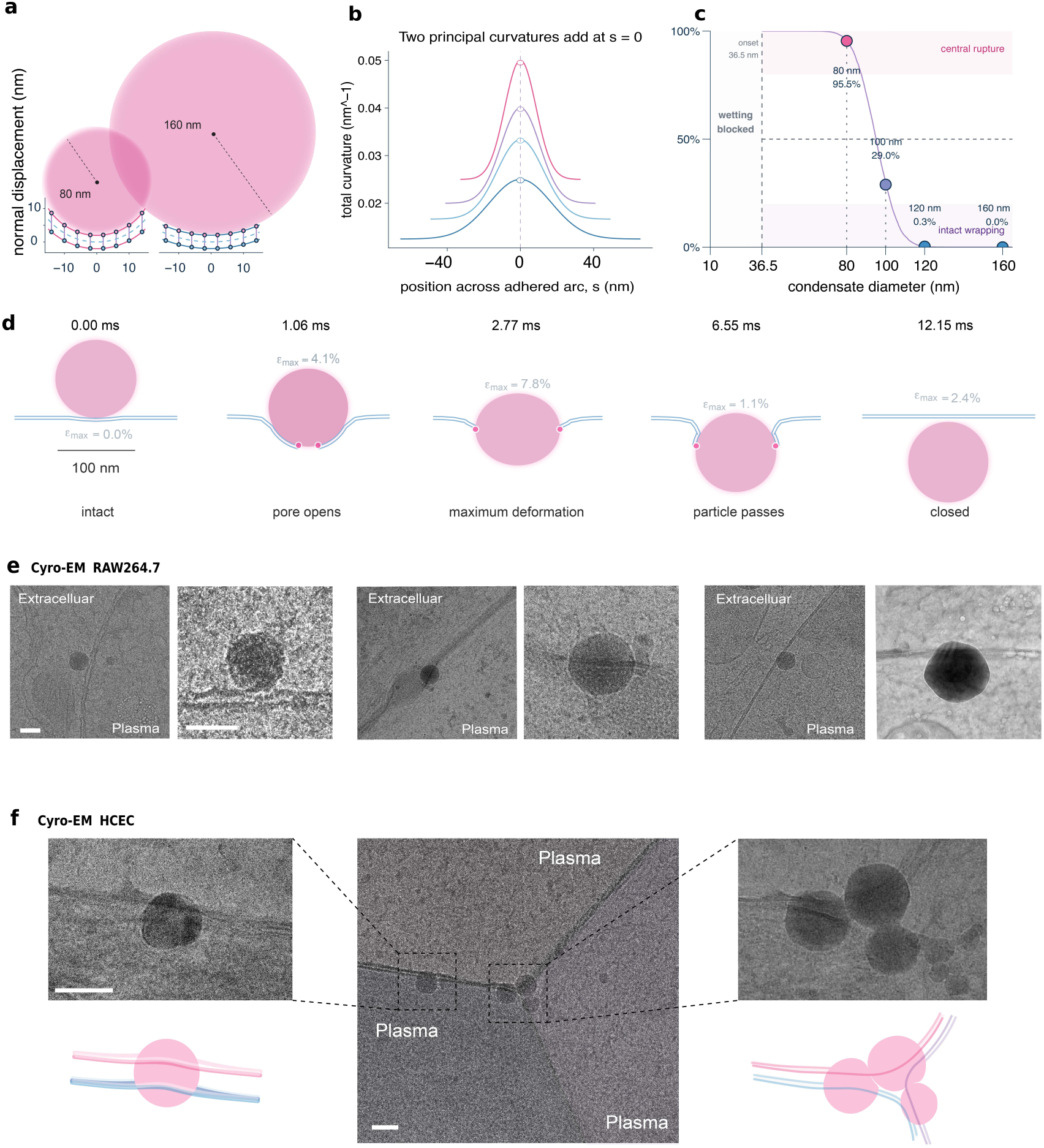
Mesoscopic properties support a transient-pore model of nanocondensate tunnelling. a, Geometrical comparison of membrane deformation beneath condensates measuring 80 or 160 nm in diameter. Curves show membrane-normal displacement across the adhered interface. **b,** Total membrane curvature, defined as the sum of the two principal curvatures, across the adhered interface for condensates measuring 80, 100, 120 or 160 nm. **c,** Experimentally constrained probability of central pore formation before 95% completion of membrane wrapping as a function of condensate diameter. Points indicate the predicted probabilities for condensates measuring 80, 100, 120 and 160 nm; shaded regions denote the central-rupture and intact-wrapping regimes. **d,** Time-resolved viscoelastic simulation of an 80-nm condensate undergoing membrane wetting, pore opening, passage and membrane closure. The maximum principal membrane strain, ε_max, is indicated at each stage. **e,** Cryo-electron micrographs and tomographic slices showing representative extracellular, membrane-associated and intracellular CPP-SF nanocondensate configurations at RAW 264.7 plasma membranes, displayed from left to right. Membrane-associated particles coincide with local bending and pore-like discontinuities, whereas intracellular particles lack a detectable continuous membrane coat. **f,** Cryo-electron tomograms of HCEC cell-cell interfaces showing nanocondensates spanning closely apposed plasma membranes. Scale bars, 100 nm (e,f).

We quantified the competition between pore formation and intact wrapping using an axisymmetric free-boundary finite-element model^36^ (Extended Data Fig. 4a-c). Explicit-lipid simulations yielded a pore-edge line tension of 5.17 pN, and the model placed the crossover between the two pathways at a particle diameter of 93.8 nm under the nominal parameter set. Incorporating the experimentally measured size distribution gave a posterior median transition diameter of 95.0 nm, with a 95% credible interval of 77.7-112.9 nm (Fig. 4c). The predicted probabilities of pore formation before complete wrapping were 95.5%, 29.0% and 0.3% for particles measuring 80, 100 and 120 nm, respectively (Fig. 4c). Particles smaller than the transition diameter therefore preferentially entered the pore-forming pathway, whereas larger particles favoured intact wrapping. Particles near 100 nm occupied a mixed-outcome regime. Conditional parameter sweeps showed that membrane tension, adhesion strength, pore-edge line tension and lipid relaxation shifted the transition diameter, with robustness analyses provided in Supplementary Data2. We next examined whether the pore-forming pathway could support particle passage and subsequent membrane closure. Viscoelastic simulations reconstructed a continuous sequence of pore opening, condensate passage and membrane resealing (Fig. 4d). A representative 80-nm particle completed this sequence within approximately 12ms, while the maximum principal strain remained below 8%. Adhesion maintained apposition between the pore rim and the condensate surface, limiting the exposed aperture during passage. Once the particle had crossed the membrane, relaxation of the wetting load allowed pore-edge line tension to restore membrane continuity.

To examine the predicted interfacial states directly, we prepared approximately 100-nm-thick cryo-FIB lamellae from RAW 264.7 cells after 30 s of SF treatment (Fig. 4e and Extended Data Fig. 4d,e). Across three independently prepared cultures, we analyzed 14 cells, 18 lamellae and 11 tomograms. Cryo-electron microscopy and tomography identified intracellular core-shell particles resembling bulk-formed CPP-SF nanocondensates. These cytoplasmic particles had a mean diameter of 104.4 ± 26.8 nm and lacked a detectable continuous membrane coat. At the plasma membrane, particle-associated regions exhibited local membrane bending and pore-like discontinuities, whereas particles located further inside the cytoplasm remained unenclosed (Fig. 4e and Extended Data Fig. 4f,g). These static ultrastructural configurations corresponded to the pore-associated and post-translocation states predicted by the model. Cryo-electron tomography of HCEC interfaces further captured nanocondensates spanning two closely apposed plasma membranes (Fig. 4f). In conjunction with the live-cell transfer events in Fig. 3f, this configuration supported direct passage across the successive donor-and recipient-cell membranes represented in Fig. 3g. Membrane perturbation remained limited under the working conditions. CPP concentrations of 1-5 μM in the presence of SF preserved cellular metabolic activity, and Annexin V and PI labelling remained limited within this concentration range (Extended Data Fig. 5a,b). Membrane damage increased at higher CPP concentrations. The concentration dependence was consistent with localized, short-lived membrane perturbation at the working concentrations, although these population-level assays did not directly measure individual pore-resealing events.

The results support a transient-pore model in which membrane affinity initiates wetting, nanoscale curvature concentrates local stress and the fluid condensate interface remains apposed to the pore rim during passage. When pore formation precedes complete wrapping, the nanocondensate crosses without acquiring a persistent membrane coat, after which pore-edge tension restores membrane continuity. This coupling of nanoscale geometry, interfacial wetting and structural reconfigurability provides a physical basis for the repeated inward and outward crossings that constitute nanocondensate tunnelling.

### Nanocondensate tunnelling supports cargo transfer across cellular and corneal barriers

Having established nanocondensate tunnelling in cellular models, we next asked whether it could operate across a native tissue barrier. The mucin-rich tear film and epithelial glycocalyx at the corneal surface provide a confined interface for in situ assembly^37,38^. We therefore applied CPP topically to associate with the glycocalyx, followed by SF to form a superficial condensate reservoir (Extended Data Fig. 6a-d). Adjusting the CPP-to-SF mass ratio (BcS) increased SF recruitment. Strong SF-Cy5.5 and CPP-FAM colocalization supported co-assembly, whereas signals in deeper epithelium suggested inward migration (Extended Data Fig. 6e).

We tracked nanocondensate distribution across four independently sampled corneas at each time point from 10 min to 1 h. At 10 min, fluorescence was concentrated in the surface reservoir, with limited extension into deeper epithelium. The surface signal subsequently declined as fluorescence increased throughout the epithelium, entered the stroma at approximately 30 min and progressed towards the endothelium. By 1 h, the reservoir was nearly depleted and fluorescence was dispersed across deeper tissue (Fig. 5a-d). Perinuclear epithelial signals were consistent with the cytosolic diffusion-like behaviour observed in cellular models, supporting stepwise intracellular migration rather than exclusively paracellular diffusion (Fig. 5b). Fluorescence was not fully transferred between successive layers, possibly because non-directional diffusion diverted particles laterally, outwards or into local retention. Thus, only a subset continued towards the endothelial side (Fig. 5e). Together, the time-resolved progression of SF signals across tightly connected corneal layers and the presence of intracellular and perinuclear puncta experimentally extended nanocondensate tunnelling from cultured cells to a native multilayered tissue barrier. Because these tissue-scale measurements did not resolve every individual membrane crossing, a paracellular contribution may accompany the observed transport. We next evaluated cargo versatility^39^. Weak-interaction-driven co-assembly enabled loading of hydrophilic and hydrophobic small molecules and biological macromolecules (Extended Data Fig. 6f and Extended Data Table 1). Adding cargo directly to SF enabled one-step formation of cargo-loaded nanocondensates on CPP-pretreated cells. Confocal imaging showed enhanced intracellular delivery of rhodamine B, siRNA and anti-VEGF antibody, each strongly colocalized with SF (Extended Data Fig. 6g-i). Quantification used four independent experiments with 30 cells per cargo and condition in each experiment; cell-level values were averaged before inference.

**Fig. 5.**
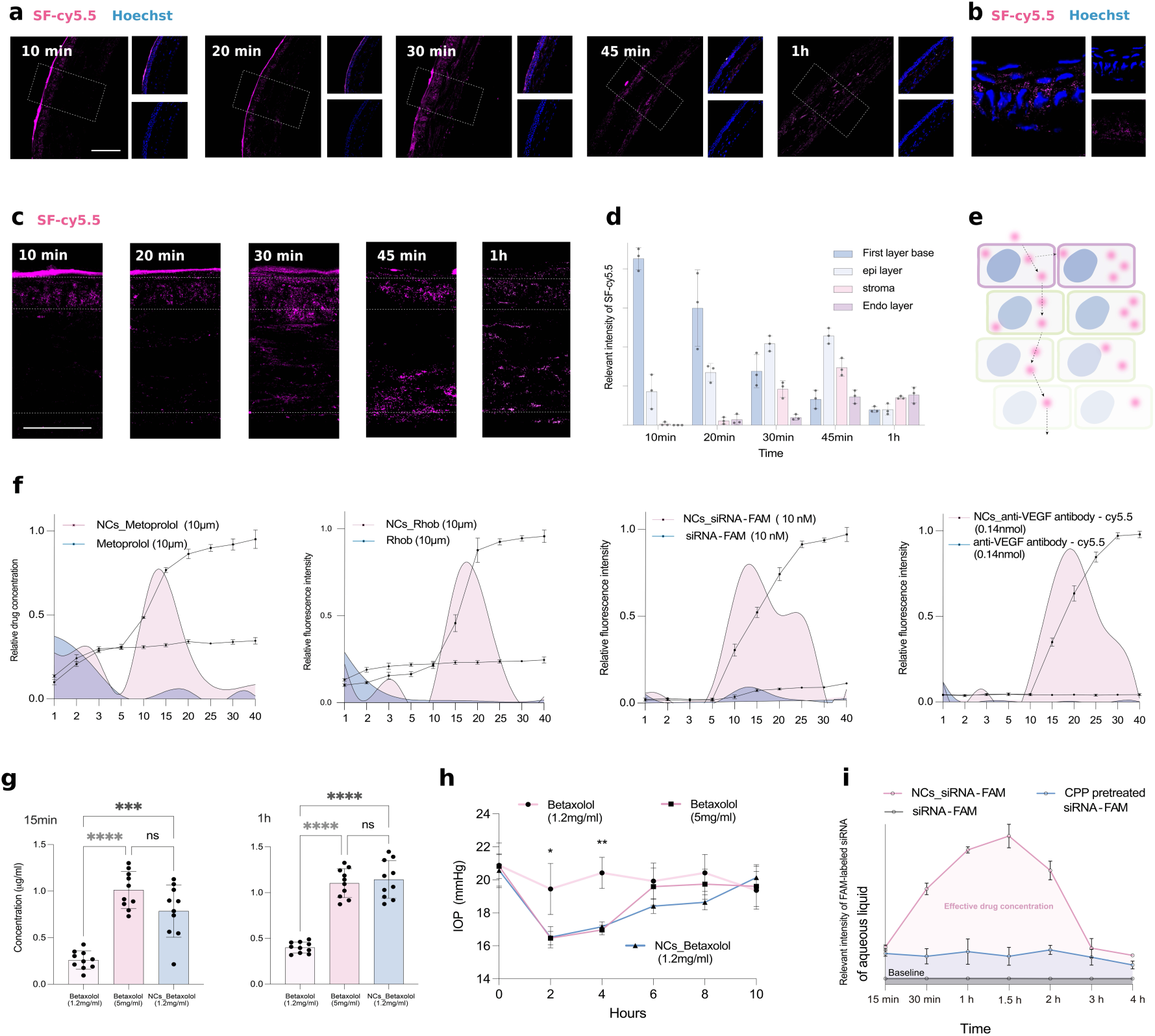
Nanocondensate tunnelling supports cargo transport across cellular and corneal barriers. a, Confocal images showing the distribution of SF-Cy5.5 across corneal tissue at the indicated times after topical administration. Boxed regions are shown at higher magnification. **b,** Higher-magnification image showing perinuclear SF-Cy5.5 signals in the corneal epithelium. **c,** Depth-resolved SF-Cy5.5 distribution from 10 min to 1 h. **d,** Quantification of SF-Cy5.5 signals in the surface reservoir, epithelium, stroma and endothelium. **e,** Model of progressive cellular transfer across successive corneal layers. **f,** Basolateral transport profiles of metoprolol, rhodamine B, FAM-labelled siRNA and Cy5.5-labelled anti-VEGF antibody across HCEC Transwell barriers, delivered as free cargoes or within nanocondensates. **g,** Betaxolol concentrations in the aqueous humour at 15 min and 1 h after topical administration of free or nanocondensate-loaded betaxolol. **h,** Intraocular pressure after treatment with free or nanocondensate-loaded betaxolol. **i,** Time course of FAM-labelled siRNA in the aqueous humour after administration of free siRNA, CPP pretreatment followed by siRNA, or siRNA-loaded nanocondensates. For a-d, n = 4 corneas from four mice per time point. For f, n = 3 independently cultured Transwell barriers; two-sided Welch’s t-tests comparing transport AUCs, with Holm correction, gave adjusted *P* = 0.014, 0.006, 0.0007 and 0.0012 for metoprolol, rhodamine B, siRNA and antibody, respectively. For g n = 10 mice per group; two-sided two-way ANOVA gave treatment-by-time interactions of *P* < 0.0001 in both experiments. For h, n = 8 mice per group; a two-sided mixed-effects model gave a treatment-by-time interaction of *P* < 0.0001. Data are mean ± s.e.m. Scale bars, 60 µm (a) and 30 µm (c).

We then established five independently cultured HCEC Transwell barriers, each maintained at a transepithelial electrical resistance of 800 ± 50 Ω·cm² (Extended Data Fig. 6j). Brief exposure to metoprolol or rhodamine B generated measurable basolateral signals, potentially through passive paracellular diffusion. Nanocondensates nevertheless increased cumulative transport and produced delayed peaks consistent with cellular uptake and re-release. Free siRNA and antibody showed little barrier crossing after a 1-min exposure, whereas nanocondensates markedly increased their basolateral signals and generated similar delayed peaks (Fig. 5f). Nanocondensates therefore transported cargoes spanning diverse physicochemical properties and molecular sizes across a tight-junction-forming cellular barrier.

Finally, we assessed corneal delivery and pharmacological efficacy in randomized, blinded mouse experiments. Aqueous-humour betaxolol and siRNA measurements used six mice per group and time point, and intraocular-pressure measurements used eight mice per group. At 15 min and 1 h, 1.2 mg ml⁻¹ betaxolol-loaded nanocondensates produced higher aqueous-humour concentrations than equal-dose free betaxolol and concentrations comparable to 5 mg ml⁻¹ free drug (Fig. 5g). The formulation produced an intraocular-pressure reduction approaching that of 5 mg ml⁻¹ free betaxolol and a more sustained effect than equal-dose free drug (Fig. 5h). FAM-siRNA-loaded nanocondensates likewise produced aqueous humour fluorescence that peaked at approximately 1.5 h and exceeded signals from free siRNA and CPP pretreatment (Fig. 5i). The modest CPP-associated signal could reflect limited transcorneal passage or interference from ocular-surface retention during aqueous humour sampling. In vivo imaging further showed stronger and more persistent ocular fluorescence after nanocondensate delivery, supporting enhanced ocular delivery and retention (Extended Data Fig. 6k,l). Thus, nanocondensate tunnelling transported small molecules and siRNA across the corneal barrier, while enabling small-molecule, siRNA and antibody transport across the tight-junction-forming HCEC barrier.

## Discussion

Here we show that in situ co-condensation of CPP and SF at the plasma membrane produces fluid nanocondensates capable of repeated inward and outward membrane crossing and intercellular transfer. These findings identify nanocondensate tunnelling as a material-state-dependent form of intercellular transport arising from a locally generated condensate state^40^. By acting as an assembly interface, the plasma membrane converts mesoscopic organization into cellular transport^21,41^.

CPP uptake has long been susceptible to artefacts introduced by chemical fixation^42^, showed that fixation can redistribute surface-bound CPP-fluorophore conjugates and produce apparent cytosolic localization. In our study, SF served as the principal transport readout, translocation events were recorded in living cells, and ultrastructural states were examined in vitrified specimens. The concordance between live-cell dynamics and cryogenic ultrastructure supports nanocondensate tunnelling as a physical translocation process.

A defining feature of nanocondensate tunnelling is that nanocondensates remain free of a persistent membrane enclosure after crossing. This unenclosed state preserves their wetting characteristics after passage and thereby their capacity to engage successive membranes. We propose that this state arises from their dual mesoscopic properties, nanoscale geometry and a fluid, reconfigurable material state. Convergent theoretical and ultrastructural evidence supports this transient-pore model, providing a testable explanation for repeated membrane crossing. Further investigation should establish how broadly this mechanism operates and identify additional processes that regulate nanocondensate tunnelling.

The functional significance of nanocondensate tunnelling lies in its capacity to carry associated material across successive cellular interfaces. Cargo experiments demonstrate transport across cellular barriers for distinct molecular classes, while corneal experiments extend tunnelling to a native multilayered tissue. Local formation of a transport-competent condensate state may inform strategies for moving biologics across restrictive barriers. Viral entry and spread^43^ offer a broader context in which analogous phase-like nanoassemblies generated by multivalent attachment and molecular clustering merit investigation. More broadly, nanocondensate tunnelling provides a testable framework for examining how mesoscopic organization enables material exchange across biological interfaces^44^.

## Methods

### Study design and biological replication

Experiments were designed as exploratory mechanistic studies. Sample sizes were specified before data collection from pilot variability; no formal power calculation was used. Unless stated otherwise, cell-based assays comprised four independent biological experiments performed on separate days with independently cultured cells and freshly prepared CPP-SF material. Cells, fields, particles and technical wells were treated as subsamples. Values were averaged within each experiment, and the independent experiment was the statistical unit. Technical replicates were not entered as independent observations. No biological replicate or animal was excluded. Prespecified technical exclusions were loss of focus, detector saturation, failed segmentation, tracking shorter than three consecutive frames, tomograms outside the specified thickness range, or control-well viability below 80% of the plate median.

### Randomization and blinding

Culture wells were assigned to treatments with a computer-generated random sequence, and treatment order was balanced across plates. Imaging fields were selected from a predefined stage-coordinate grid after random shuffling. The operator preparing CPP-SF formulations was not blinded because concentrations were visible during dosing. Acquisition settings were fixed before imaging, files were assigned anonymous codes, and fluorescence quantification, particle sizing, FRAP fitting, trajectory detection, tomogram classification, flow-cytometry gating, tissue-intensity measurement, aqueous-humour assays and intraocular-pressure analysis were performed without knowledge of treatment. Codes were broken after the analysis table had been locked.

### Definition and calculation of binary-component stoichiometry

Binary-component stoichiometry (BcS) denotes the feed mass ratio of cell-penetrating peptide (CPP) to silk fibroin (SF). A mass ratio was used rather than a molar ratio because the CPP variants differed in sequence and molecular mass. BcS is reported as 1:n, where the CPP mass is normalized to 1; thus, BcS = 1:14 indicates one mass unit of CPP per 14 mass units of SF.

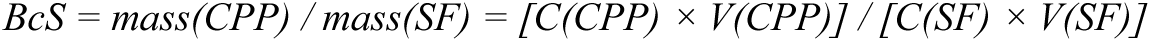

For bulk co-assembly, BcS was calculated from the masses mixed in solution. For cell-surface and ocular-surface assembly, BcS is the nominal feed ratio calculated from the amount of CPP applied during preloading and the amount of SF applied during the second step. Because washing removes unbound CPP, the local interfacial ratio can differ from the nominal value. Unless stated otherwise, BcS values for cellular and ocular experiments therefore denote nominal feed ratios. Low-to-intermediate BcS conditions that produced nanoscale particles, fluorescence recovery and limited irreversible precipitation were classified as liquid-biased. Higher BcS conditions that produced broad size distributions, fibrils or centrifugation-resistant aggregates were classified as aggregate-biased.

### Preparation of silk fibroin

Aqueous SF was prepared from Bombyx mori cocoons by degumming and lithium bromide dissolution. Cocoons were cut into pieces and boiled in 0.02 M Na₂CO₃ for 30 min to remove sericin. Fibres were rinsed with ultrapure water until the wash was clear, dried overnight at room temperature and dissolved in 9.3 M LiBr for 4 h at 60 °C. The solution was dialysed against ultrapure water for 48-72 h with at least six water changes, clarified at 9,000g for 20 min and passed through a 0.22-µm filter. SF concentration was determined gravimetrically from dried aliquots. Stock solution was stored at 4 °C and used within 7 d. Unless stated otherwise, final SF concentrations were 4 mg ml⁻¹ for bulk experiments, 0.5 mg ml⁻¹ for cellular and Transwell experiments, and 1 mg ml⁻¹ for ocular-surface experiments.

### Preparation and fluorescent labelling of CPP and SF

CPP and tandem-duplicated CPP (dCPP) were dissolved in ultrapure water or PBS, aliquoted and stored at −80 °C. Stocks were thawed once and diluted immediately before use. SF concentration was held constant within each experiment, and CPP concentration was varied to set BcS. Fluorophore-conjugated CPPs carried FAM, Cy3 or Cy5 as indicated. For SF labelling, SF was reacted with an NHS-activated dye in 0.1 M bicarbonate buffer, pH 8.3, for 2 h in the dark. Free dye was removed by dialysis or centrifugal ultrafiltration until filtrate fluorescence reached background. Labelled material was mixed with unlabelled material at a fixed fraction within each experiment to minimize dye-dependent changes in assembly. All labelled reagents were protected from light.

### Bulk CPP-SF co-assembly and phase-state classification

SF was diluted to 4 mg ml⁻¹, and CPP was added dropwise under gentle pipette mixing to generate the indicated BcS values. Mixtures were incubated for 1-5 min at room temperature and analysed immediately. Turbidity was measured at 600 nm in transparent 96-well plates. For phase diagrams, SF and CPP concentrations were varied systematically, and each sample was classified from turbidity, microscopic morphology, particle-size distribution and centrifugation-resistant material. Phase boundaries were reproduced in three independent preparation series. Quantitative turbidity measurements comprised four independently prepared samples per condition.

### Centrifugation-resistant precipitate assay

Freshly assembled CPP-SF samples were centrifuged at 20,000g for 10 min at room temperature. The supernatant was removed, and the pellet was washed once with ultrapure water, dried to constant mass and weighed. The precipitated fraction was calculated relative to the total CPP and SF input mass. Four independent assembly preparations were analysed; any repeat weighing of the same pellet was treated as a technical measurement and averaged before analysis.

### Dynamic light scattering

Hydrodynamic size distributions were measured at 25 °C immediately after assembly. Samples were diluted only when the instrument count rate exceeded the validated range. Three consecutive measurements were acquired from each preparation and averaged. Three independent preparations were analysed per BcS condition, and the preparation mean was the statistical unit.

### Acetone desolvation

For comparison of hydrated and collapsed structures, acetone was added slowly to freshly assembled CPP-SF nanocondensates under gentle stirring until particle solidification was complete. Particles were collected at 20,000g for 10 min, washed twice with ultrapure water and resuspended for TEM and size analysis. Desolvated particles were used only as a structural comparator and were not used in cellular or ocular delivery experiments.

### Transmission electron microscopy and cryo-TEM

For negative-stain TEM, 5 µl of freshly prepared sample was adsorbed onto a glow-discharged carbon-coated copper grid for 1 min, blotted and stained twice with 2% uranyl acetate. For cryo-TEM, 3 µl of sample was applied to a glow-discharged holey-carbon grid, blotted under controlled humidity and vitrified in liquid ethane. Grids were stored under liquid nitrogen until imaging. Images were acquired at fixed magnification within each comparison. Particle diameter, core diameter and shell thickness were measured from coded images with a prespecified segmentation workflow. Only particles with a complete, in-focus boundary were included. Extended Data Fig. 2d used 60 particles per group sampled equally from three independent preparations; preparation-level means were used for inference.

### FRET assay

CPP and SF were labelled as donor and acceptor, respectively; the reciprocal configuration was used as a confirmatory control. Components were mixed at the indicated BcS and spectra were collected after assembly using donor excitation. Donor-only and acceptor-only controls were used to correct bleed-through and direct excitation. Corrected acceptor emission was normalized to donor signal within each preparation.

### Spectroscopy and zeta-potential measurements

For Fourier-transform infrared spectroscopy, assemblies were lyophilized and spectra were acquired over the amide-I region. Component bands representing β-sheet, β-turn and random-coil structures were fitted with the same constraints for all samples. Circular-dichroism spectra of CPP were recorded before and after addition of NaCl, FeCl₃ or CaCl₂, and changes in the α-helical signal were quantified. Raman spectra were baseline-corrected using a fixed pipeline, and protonated amino-group bands were compared across conditions. Zeta potential was measured at 25 °C in four independent preparations, each represented by the mean of three technical measurements. Spectroscopy experiments were reproduced in three independent preparations.

### STED microscopy

Fluorescently labelled CPP and SF were assembled in solution or on cell membranes and imaged by sequential-channel stimulated-emission-depletion microscopy. Laser power, detector gain, depletion power and reconstruction settings were fixed within each experiment. For bulk particles, radial intensity profiles were aligned to the particle boundary to compare CPP-enriched shells with SF-enriched cores. For membrane-associated assemblies, punctum size and morphology were quantified from coded images using a fixed threshold.

### FRAP

For bulk FRAP, labelled condensates were transferred to glass-bottom dishes and a defined region within one condensate was bleached. For cell-surface FRAP, cells were exposed to CPP for 30 s, washed once and treated with 0.5 mg ml⁻¹ SF at 4 °C to promote the accumulation and retention of condensates at the plasma membrane. An arc-shaped membrane region of interest (ROI), with a curvature corresponding to a 5-μm-diameter circle and a radial width of 0.5 μm, was then photobleached. Following bleaching, the temperature was gradually increased from 4 °C to room temperature, and 0.1 mg ml⁻¹ SF was added as the recovery solution. Fluorescence recovery was recorded for 190 s. Background-subtracted fluorescence intensity was corrected using an unbleached control region and normalized to the pre-bleach intensity. Mobile fractions and recovery curves were calculated using the same model for all conditions. Eight cells per condition were analysed across four independent experiments, with measurements from individual cells averaged within each experiment before statistical inference.

### Cell culture

RAW 264.7 and BV2 cells were maintained in high-glucose DMEM, and HCEC, RPE and SRA cells were maintained in DMEM/F12. Media contained 10% fetal bovine serum and 1% penicillin-streptomycin. Cells were cultured at 37 °C with 5% CO₂, used at 60-80% confluence and discarded after 20 passages. Cell stocks were authenticated before the study, and cultures were confirmed to be mycoplasma-free. For imaging, cells were plated on glass-bottom dishes 18-24 h before treatment. Residual serum was removed with serum-free medium immediately before CPP exposure. Dil or FM1-43 was used to visualize the plasma membrane, and Hoechst 33342 was used when nuclear counterstaining was required.

### In situ assembly of CPP-SF nanocondensates on cells

Cells were incubated with 1-5 µM CPP for 30 s at 37 °C. The CPP solution was removed, and cells were washed once with serum-free medium to remove unbound peptide while retaining membrane-associated CPP. SF at 0.5 mg ml⁻¹ was then added for 1 min to induce co-condensation at the plasma membrane. The working BcS was adjusted by changing CPP concentration while keeping SF constant. Cells were imaged immediately under live-cell conditions or fixed in 4% paraformaldehyde for 10 min. The same nominal BcS was tested first across cell types; CPP loading was then adjusted to compensate for cell-type-dependent surface adsorption.

### Confocal imaging and quantification of cellular entry

CPP, SF, cargo, membrane and nuclear channels were acquired sequentially to minimize spectral bleed-through. Time-course images were collected at 30 s, 1, 2, 5, 10, 15, 30 and 40 min after SF addition as required. Laser power, detector gain and thresholds were fixed within each experiment. Membrane and intracellular masks were generated using a fixed automated pipeline from membrane, nuclear and transmitted-light images and then verified while blinded. The intracellular fraction was the SF signal within the cell-body mask divided by total cell-associated SF signal. Colocalization was quantified using Pearson correlation and line-scan profiles. For Fig. 2c,f, Fig. 3a and Extended Data Fig. 3d-i, 30 cells per condition were sampled in each of four independent experiments and averaged within experiment. Particle sizing used 186 particles from 62 cells in three independent experiments; 147 ± 67.4 nm is the particle-level mean ± s.d., whereas statistical inference used experiment-level means.

### Live-cell event detection and particle tracking

Time-lapse sequences were acquired at 10 frames s⁻¹ immediately after SF addition. Entry, outward translocation or intercellular transfer was scored only when a punctum crossed a segmented membrane boundary and remained on the opposite side for at least three consecutive frames. Trajectories shorter than three frames, merged puncta and saturated signals were excluded by prespecified criteria. Blinded analysis identified 47 entry events in 31 cells, 22 outward-translocation events in 18 cells and 12 complete transfers across eight cell pairs, each sampled over four independent experiments. Intracellular diffusion analysis used 30 particles from 10 cells across three independent experiments.

### Cryogenic electron microscopy and tomography of cells

RAW 264.7 cells treated with CPP-SF nanocondensates were vitrified immediately after the indicated incubation. Surface morphology was first examined under cryogenic conditions. Candidate cells were selected by systematic sampling of every third eligible grid square after a random starting position. Cryo-focused ion-beam milling generated approximately 100-nm lamellae, which were transferred under cryogenic conditions for electron microscopy and tomography. Intracellular particles were identified from size, morphology and core-shell architecture; the presence of a continuous surrounding membrane and local pore-like openings was scored independently. Three preparations yielded 18 lamellae from 14 cells; 11 tomograms met the prespecified thickness and alignment criteria. Particle and membrane states were classified from coded tomograms by two blinded observers, with disagreements resolved by consensus. Seven closely apposed cell-cell interfaces were analyzed in six tomograms. Representative images were chosen only after the coded set had been scored.

### Cell viability, flow cytometry and lysosomal colocalization

For CCK-8 measurements, cells received CPP alone, SF alone or sequential CPP and SF treatment under the delivery conditions. Metabolic activity was measured after addition of CCK-8 reagent and normalized to untreated cells. Five independent experiments were performed, each represented by the mean of four technical wells per condition. For Annexin V and propidium iodide analysis, cells were washed, stained and analyzed by flow cytometry. Four independent experiments were performed with at least 20,000 singlets per sample. Debris, doublets and non-cellular events were removed by a fixed sequential gate; Annexin V and propidium iodide gates were set from unstained and single-stained controls before coded samples were analyzed. For lysosomal colocalization, cells were stained with LysoTracker and imaged by sequential-channel confocal microscopy. Colocalization and trajectory analyses in Extended Data Fig. 5c,d used 30 particles from 10 cells across three independent experiments.

### Cargo loading and loading calculations

Cargo was mixed directly with SF before the second assembly step. Hydrophilic tracers, siRNA and proteins were dissolved in SF solution. Hydrophobic small molecules were first prepared as concentrated stocks and then diluted into SF under gentle mixing. SF was used at 4 mg ml⁻¹ for bulk loading, 0.5 mg ml⁻¹ for cellular and Transwell delivery, and 1 mg ml⁻¹ for ocular delivery. Free cargo and CPP-pretreated cargo without SF were included as controls. For bulk loading measurements, unencapsulated cargo was removed by dialysis or centrifugal ultrafiltration. Fluorescent cargo was quantified against an external fluorescence standard curve, and small-molecule drug was quantified by LC-MS.

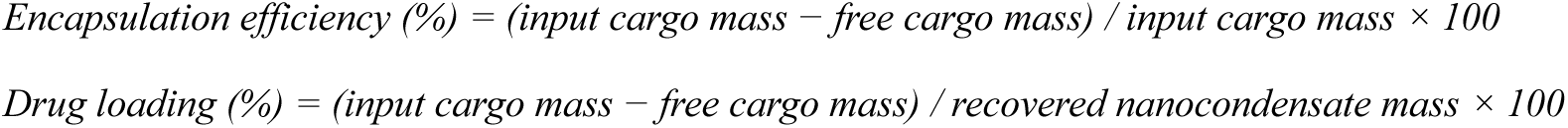

### Intracellular cargo delivery

Cells were exposed to CPP for 30 s, washed once and treated for 1 min with cargo-containing SF at 0.5 mg ml⁻¹. Extracellular cargo was removed by washing before immediate or delayed imaging. Rhodamine B, FAM-labelled siRNA and fluorescent anti-VEGF antibody were analysed in parallel with free-cargo and CPP-only controls. Intracellular fluorescence and colocalization with SF were quantified from coded confocal images. Four independent experiments were performed, with 30 cells per cargo and condition averaged within each experiment.

### Transwell epithelial transport

HCECs were seeded on Matrigel-coated Transwell inserts and cultured until a confluent barrier formed. Transepithelial electrical resistance (TEER) was measured before and after exposure; only inserts with stable baseline values of 800 ± 50 Ω·cm² were included. CPP was applied apically for 30 s and removed, after which cargo-containing SF at 0.5 mg ml⁻¹ was applied for 1 min. The apical compartment was then washed. Basolateral samples were collected at prespecified times and replaced with equal volumes of fresh medium. Metoprolol was quantified by LC-MS; rhodamine B, FAM-labelled siRNA and Cy5.5-labelled anti-VEGF antibody were quantified against fluorescence standard curves. Five inserts generated from independent cultures were used per cargo and condition. Cumulative transport and area under the concentration-time curve were calculated for each insert, which was the statistical unit. Sample order was randomized, and the analyst was blinded. Post-treatment TEER was used to verify that transport was not attributable to gross barrier disruption.

### Animals and ethical approval

C57BL/6J mice aged 8-10 weeks were maintained under a 12-h light-dark cycle with food and water available ad libitum. Experiments were approved by the Capital Medical University Animal Ethics Committee (protocol AEEI-2026-127). Sex-balanced blocks were randomized by body weight to treatment, with three females and three males in groups of six and four females and four males in groups of eight. One eye per mouse was studied; the contralateral eye was not treated as an independent control. Corneal distribution used four mice per treatment and time point. Aqueous-humour betaxolol and siRNA measurements used six mice per group and time point. Intraocular-pressure studies used eight mice per group, and in vivo fluorescence used six mice per group. The dosing operator was aware of formulation; imaging, pressure measurement, sample quantification and analysis were blinded.

### Topical ocular assembly and corneal imaging

Mice were anaesthetized for topical administration. A 5-µl CPP solution was placed on the central cornea for 30 s, and unbound solution was removed without touching the epithelium. A 5-µl SF solution at 1 mg ml⁻¹, with cargo where indicated, was then applied to the same surface to induce interfacial co-condensation. CPP concentration was varied to set nominal BcS. At 10, 20, 30 or 60 min, eyes were washed with PBS, collected, embedded in optimal-cutting-temperature compound and cryosectioned through the central cornea. Confocal images were acquired with fixed settings. Surface reservoir, epithelium, stroma and endothelium were defined from nuclear staining and tissue morphology, and fluorescence was quantified with a blinded fixed-mask pipeline. Four mice contributed one cornea each per treatment and time point; the cornea was the statistical unit.

### In vivo fluorescence imaging of ocular siRNA

Mice received topical free FAM-labelled siRNA, CPP-pretreated FAM-labelled siRNA without SF, or CPP-SF nanocondensates containing FAM-labelled siRNA. Ocular fluorescence was acquired at the prespecified time points with identical illumination, exposure and analysis regions. Background-subtracted signal was normalized to the first post-dose image for kinetic analysis. Six mice were used per group. Animal order was randomized at acquisition, and group identity remained masked during quantification.

### Aqueous-humour sampling and cargo quantification

At the indicated time points after topical administration, the ocular surface was washed thoroughly to remove residual formulation. Aqueous humour was released by corneal puncture and recovered by irrigation with a fixed volume of PBS. Betaxolol was quantified by LC-MS using matrix-matched calibration standards and an internal standard. FAM-labelled siRNA was quantified against a matrix-matched fluorescence standard curve. Six mice were analysed per treatment and time point; each mouse contributed one sample and was the statistical unit. Sample identifiers were coded before instrument analysis.

### Intraocular-pressure measurement

Intraocular pressure was measured before dosing and at the indicated post-dose time points with a rebound tonometer. Mice were measured in randomized order by an operator unaware of treatment. Six valid consecutive readings were averaged for each eye at each time point. Eight mice were used per treatment; the mouse was the statistical unit. Betaxolol-loaded CPP-SF nanocondensates at 1.2 mg ml⁻¹ were compared with equal-dose free drug and 5 mg ml⁻¹ free drug.

### Droplet-scale theory and state-map calculations

Calculations were implemented in custom Python code. Equal-droplet radii from 50 nm to 2 µm were evaluated. Brownian encounter frequency was scaled as R⁻³, capillary free-energy release as −R² and the electrostatic approach barrier as Rψ², where R is droplet radius and ψ is surface potential. A dimensionless state map combined non-electrostatic cohesion, charge-dependent repulsion and network solidification to identify dispersed, nanocondensate, coarsening and gelled regimes. Deterministic model curves were not treated as biological replicates, and no null-hypothesis test was applied.

### Membrane-mechanics and molecular-simulation analyses

An axisymmetric free-boundary model represented a finite-thickness bilayer interacting with a spherical condensate. Bending, in-plane tension, adhesion, leaflet-area mismatch and pore-edge line tension were varied over the ranges specified in the computational design. Pore-before-wrap was recorded when a central pore became energetically accessible before 95% geometric wrapping. The nominal crossover diameter was estimated from the intersection of the pore and intact-wrapping branches. Conditional sensitivity analysis comprised 336 parameter sets and 128 directional branches. Explicit-lipid restrained-pore simulations comprised 24 independent trajectories. Pore-edge line tension was fitted from the restrained-pore free-energy profile, and its 95% interval was obtained from 20,000 bootstrap resamples. These trajectories and parameter branches are computational sampling units, not biological replicates.

The experimentally measured particle-size distribution was combined with the membrane-model transition curve to estimate the probability of pore formation as a function of particle diameter. Weakly informative priors were used for the crossover position and slope; posterior summaries were calculated after convergence of four independent chains. Viscoelastic simulations then propagated membrane deformation, pore opening, particle passage and resealing through time. Model outputs were compared with live-cell event duration and cryo-electron-microscopy states, but experimental observations were not used as replicate observations for computational parameter sweeps.

### Statistical analysis

Analyses were performed in R 4.4.1 and GraphPad Prism 10.2. Tests were two-sided. Normality of experiment-level residuals was assessed with the Shapiro-Wilk test and variance homogeneity with Levene’s test. Two groups were compared with an unpaired Welch t-test. Three or more groups were compared by one-way ANOVA followed by Holm-Šidák-adjusted comparisons. Factorial experiments used two-way ANOVA. Time courses and repeated ocular measurements used mixed-effects models with experiment or animal as a random intercept and the Geisser-Greenhouse correction. Holm-Šidák adjustment was applied within each prespecified family of comparisons. For the four Transwell cargo AUC comparisons, Holm adjustment was applied across cargoes. When assumptions were not met, Kruskal-Wallis tests with Dunn-adjusted comparisons were used. Data are mean ± s.e.m. unless otherwise stated; particle-size distributions and bulk physicochemical measurements are mean ± s.d. where indicated. Adjusted P < 0.05 was considered significant, and confidence intervals are 95%.

