## Extended Data for "Nanocondensate tunnelling"

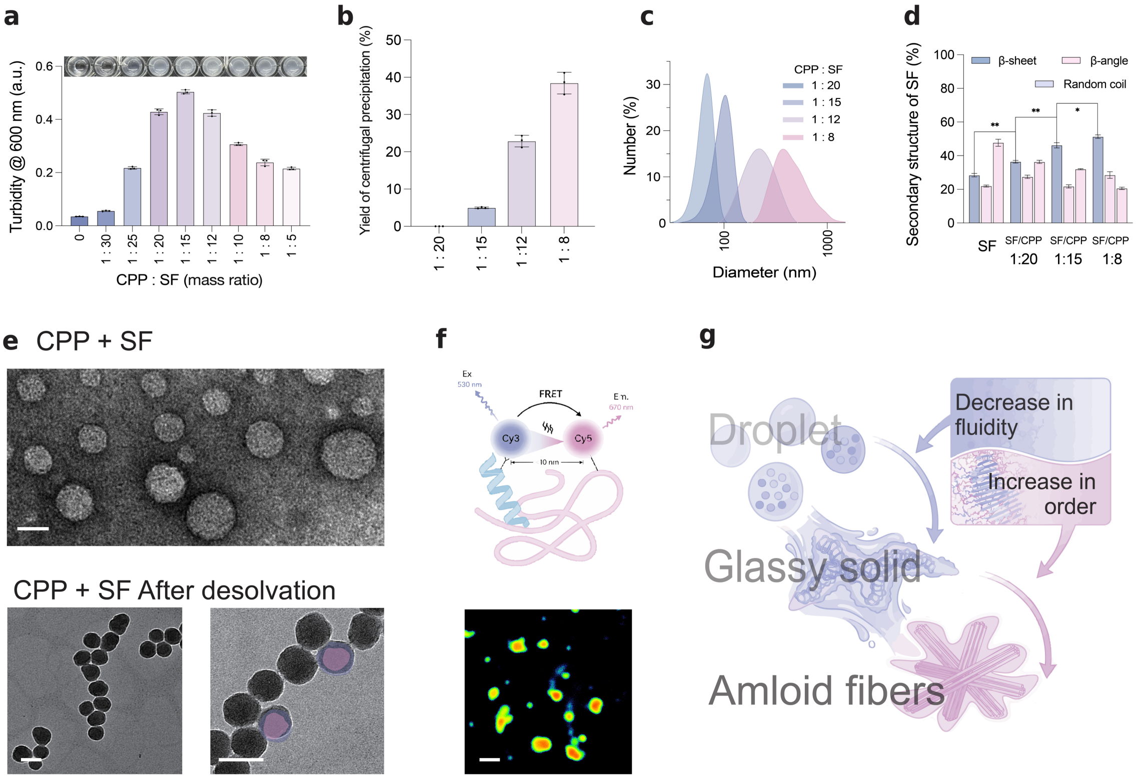


**Extended Data Fig. 1 | BcS controls the material state of CPP-SF assemblies.**

**a,** Turbidity at 600 nm and representative photographs of CPP-SF mixtures at the indicated CPP-to-SF mass ratios. **b,** Fraction of material recovered by centrifugation. **c,** Number-weighted particle-diameter distributions. **d,** SF β-sheet, β-angle and random-coil contents derived from amide-I spectral decomposition. **e,** Electron micrographs before and after desolvation; false colour delineates the core and shell. **f,** Cy3-Cy5 FRET scheme and representative FRET map of labelled CPP and SF. **g,** Proposed BcS-dependent progression from fluid nanocondensates to solid-like and fibrillar assemblies. For a and b, n = 4 independent assembly preparations. For c and d, n = 3 independent preparations, each measured in technical triplicate and averaged before analysis. FRET imaging in f was reproduced in four independent preparations. Bars in a,b,d show mean ± s.d. Group comparisons in a,b,d used two-sided one-way ANOVA with Holm-Šidák correction. Exact adjusted *P* values are provided in Source Data. Scale bars, 100 nm (e,f).

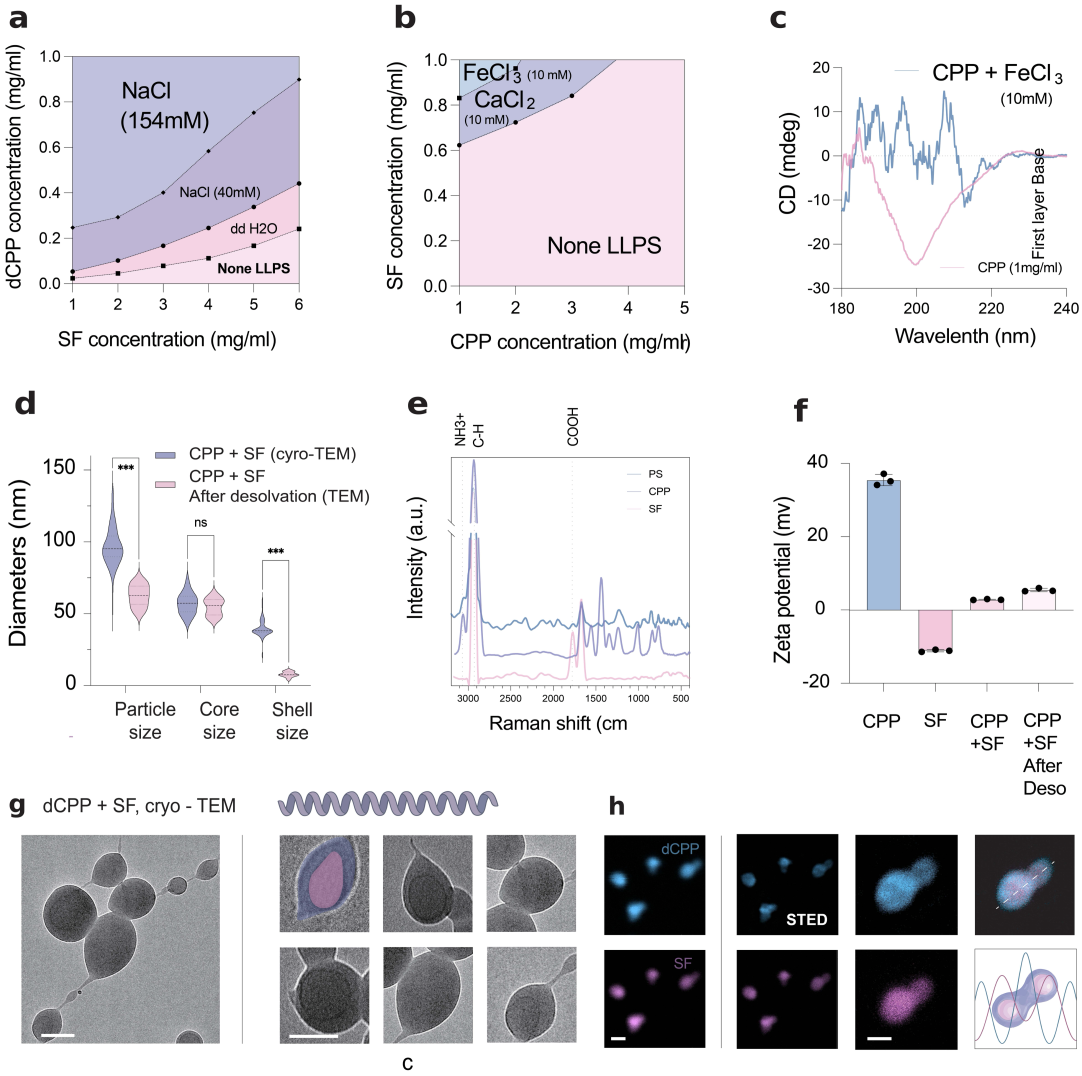


**Extended Data Fig. 2 | Charge organization and hydration govern CPP-SF nanocondensate formation.**

**a,** Phase diagram of dCPP and SF at the indicated NaCl concentrations. **b,** CPP-SF phase boundaries in 10 mM $FeCl_3$ or $CaCl_2$. **c,** Circular-dichroism spectra of CPP with or without 10 mM $FeCl_3$. **d,** Particle, core and shell diameters under hydrated cryo-TEM conditions and after desolvation. **e,** Raman spectra of phase-separated CPP-SF assemblies (PS), CPP and SF. **f,** ζ-potentials of CPP, SF and CPP-SF assemblies before and after desolvation. **g,** Cryo-TEM images of dCPP-SF nanocondensates. **h,** Conventional and STED microscopy showing peripheral dCPP enrichment and an SF signal extending across the condensate and peaking towards its centre; right, radial intensity profiles. Phase diagrams and spectra in a-c,e were reproduced in three independent preparations. In d, $n = 60$ particles per condition, sampled equally from three independent preparations; violins show particle-level distributions, whereas preparation means were the independent units for inference. In f, $n = 4$ independent preparations; points denote preparations and bars show mean ± s.e.m. Images in g,h are representative of three independent preparations. Comparisons in d,f used two-sided one-way ANOVA with Holm-Šidák correction. Exact adjusted *P* values are provided in Source Data. Scale bars, 100 nm (g,h).

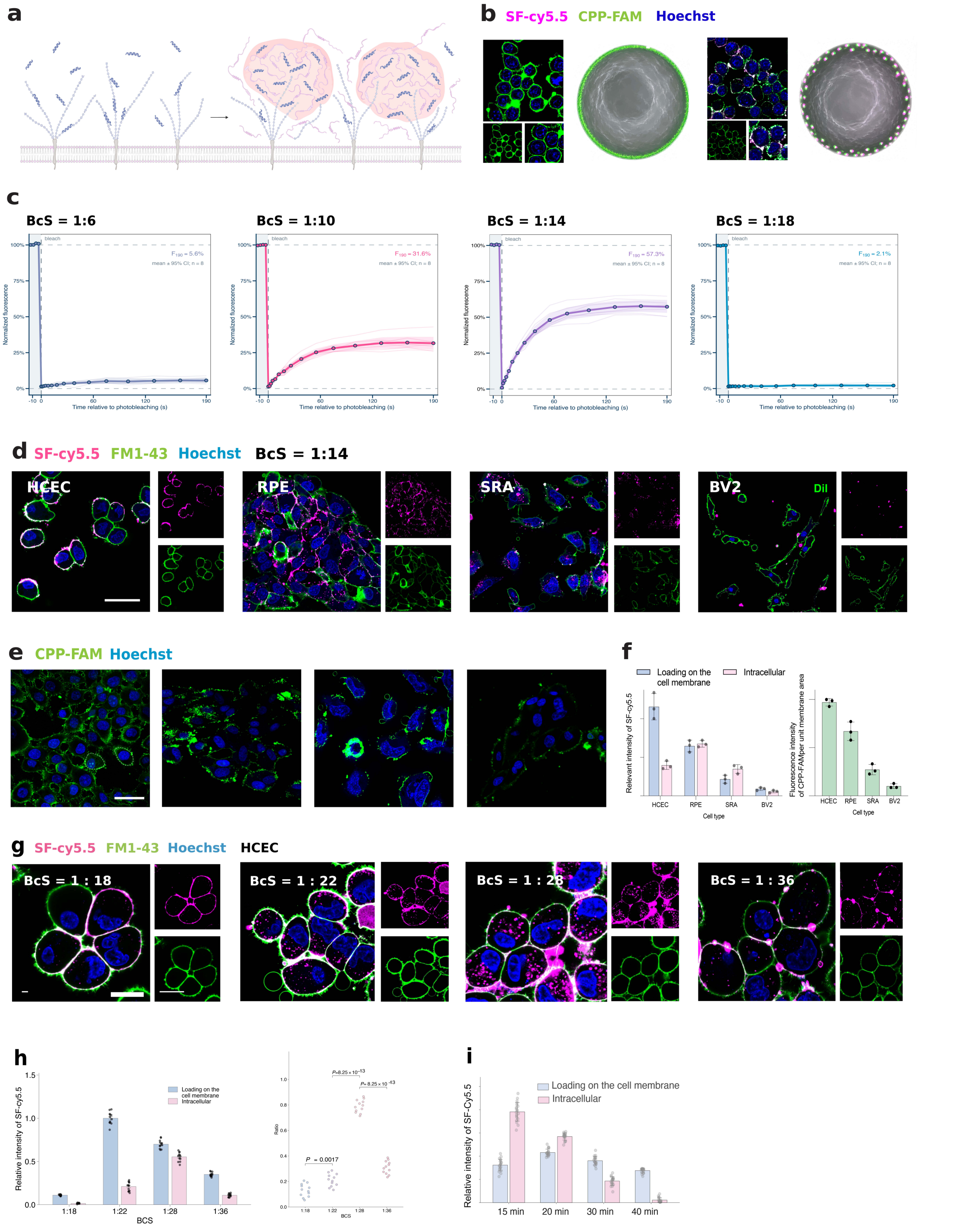

**Extended Data Fig. 3 | Surface CPP loading and condensate fluidity define the BcS window for cellular entry.** a,b, Surface CPP adsorption and membrane-confined CPP-SF co-assembly. c, FRAP curves; recovery at 190 s was 5.6%, 31.6%, 57.3% and 2.1% at BcS = 1:6, 1:10, 1:14 and 1:18 (n = 8 cells per condition across four experiments; shading, 95% confidence interval; one-way ANOVA, Holm-Šidák correction). d-f, SF distribution and CPP loading across cell types. g,h, BcS-dependent HCEC distribution and quantification. i, Time-dependent intracellular-to-membrane redistribution. For d-i, n = 4 experiments with 20 cells averaged per condition or time point; Holm-Šidák-adjusted tests and exact P values are shown. Scale bars, 10 μm (b,d,e,g).

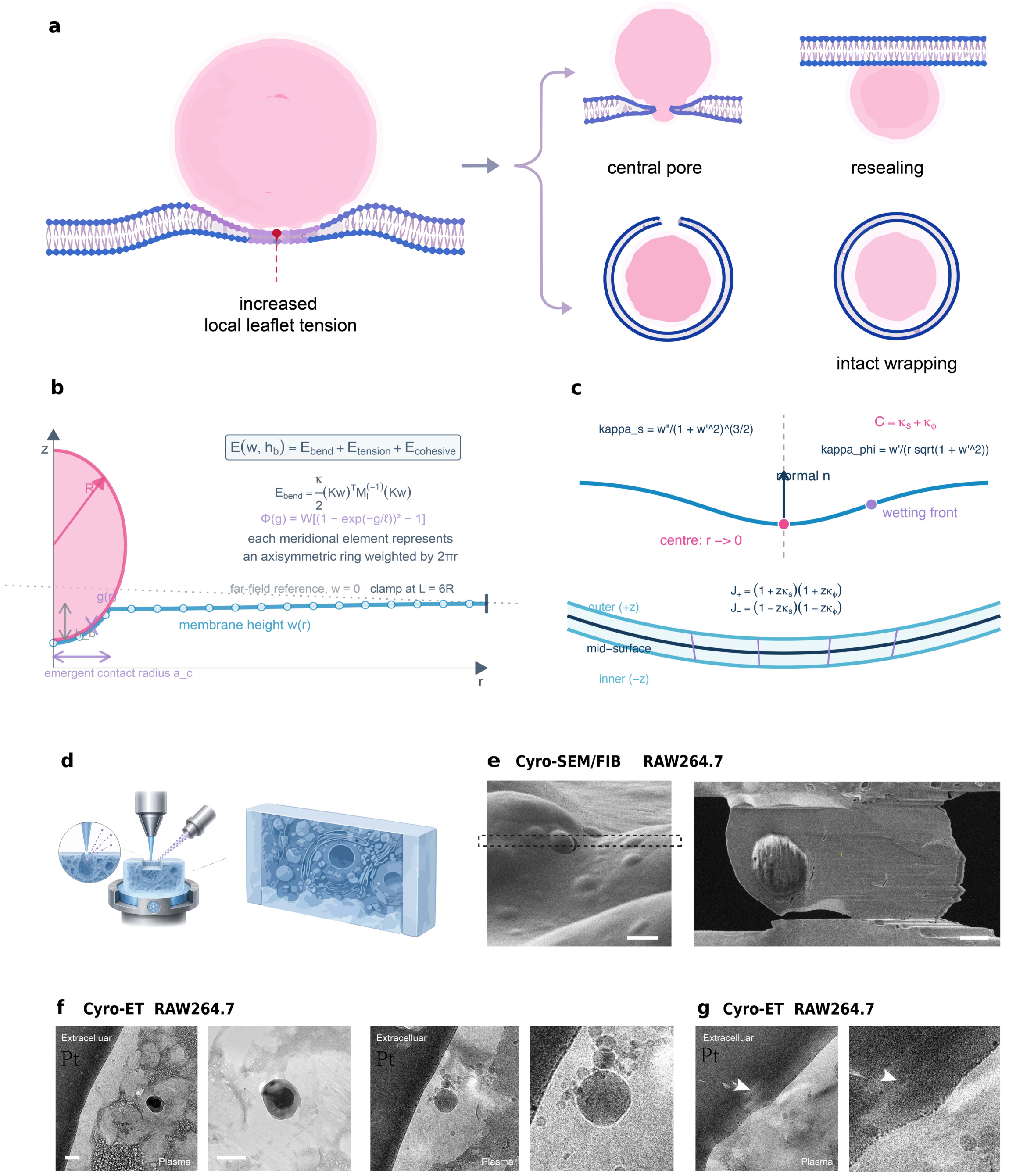


**Extended Data Fig. 4 | Membrane mechanics and ultrastructural validation of transient-pore-mediated nanocondensate translocation.**

a, Competing central-pore and intact-wrapping pathways. b,c, Axisymmetric free-boundary model geometry and curvature definitions for a finite-thickness bilayer. d, Cryo-FIB workflow. e, Cryo-SEM/FIB lamella preparation in RAW 264.7 cells. f,g, Tomographic states of contact, pore-like opening and passage across a single RAW 264.7 plasma membrane. Panels d-g summarize three independent preparations comprising 14 cells, 18 lamellae and 11 tomograms. Computational panels are not biological replicates, and no null-hypothesis test was applied. Scale bars, 5 μm (e) and 100 nm (f,g).

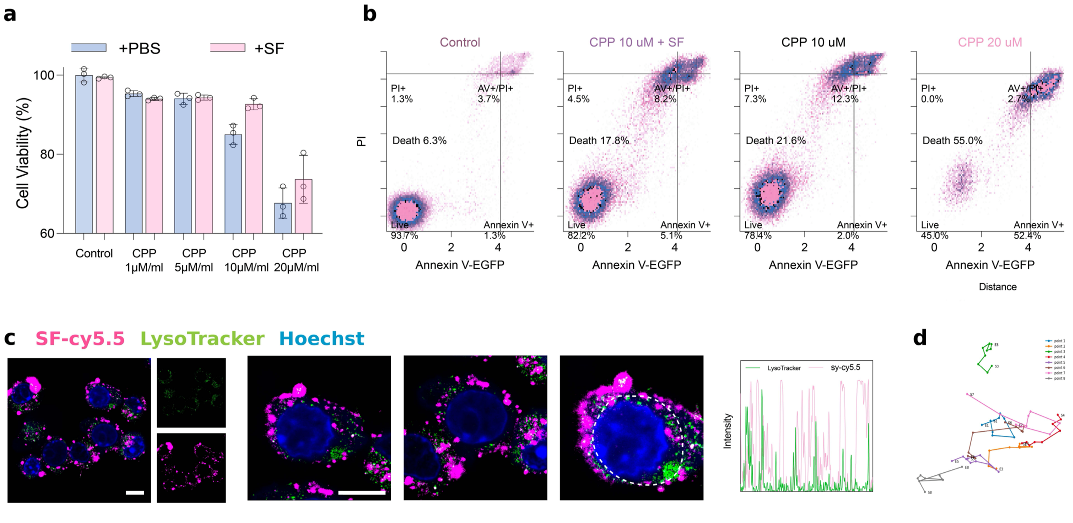


**Extended Data Fig. 5 | Cellular effects and intracellular fate of CPP-SF nanocondensates.**

a, CCK-8 viability. b, Annexin V-EGFP/PI flow cytometry. c, SF-Cy5.5, LysoTracker and Hoechst signals. d, Intracellular trajectories. In a, n = 5 independent experiments, each the mean of four technical wells. In b, n = 4 independent experiments with at least 20,000 singlets per sample. Viability and flow data are mean ± s.e.m. and used two-sided two-way ANOVA with Holm-Šidák correction. Panels c and d summarize 30 particles from 10 cells across three experiments. Scale bars, 5 µm (c).

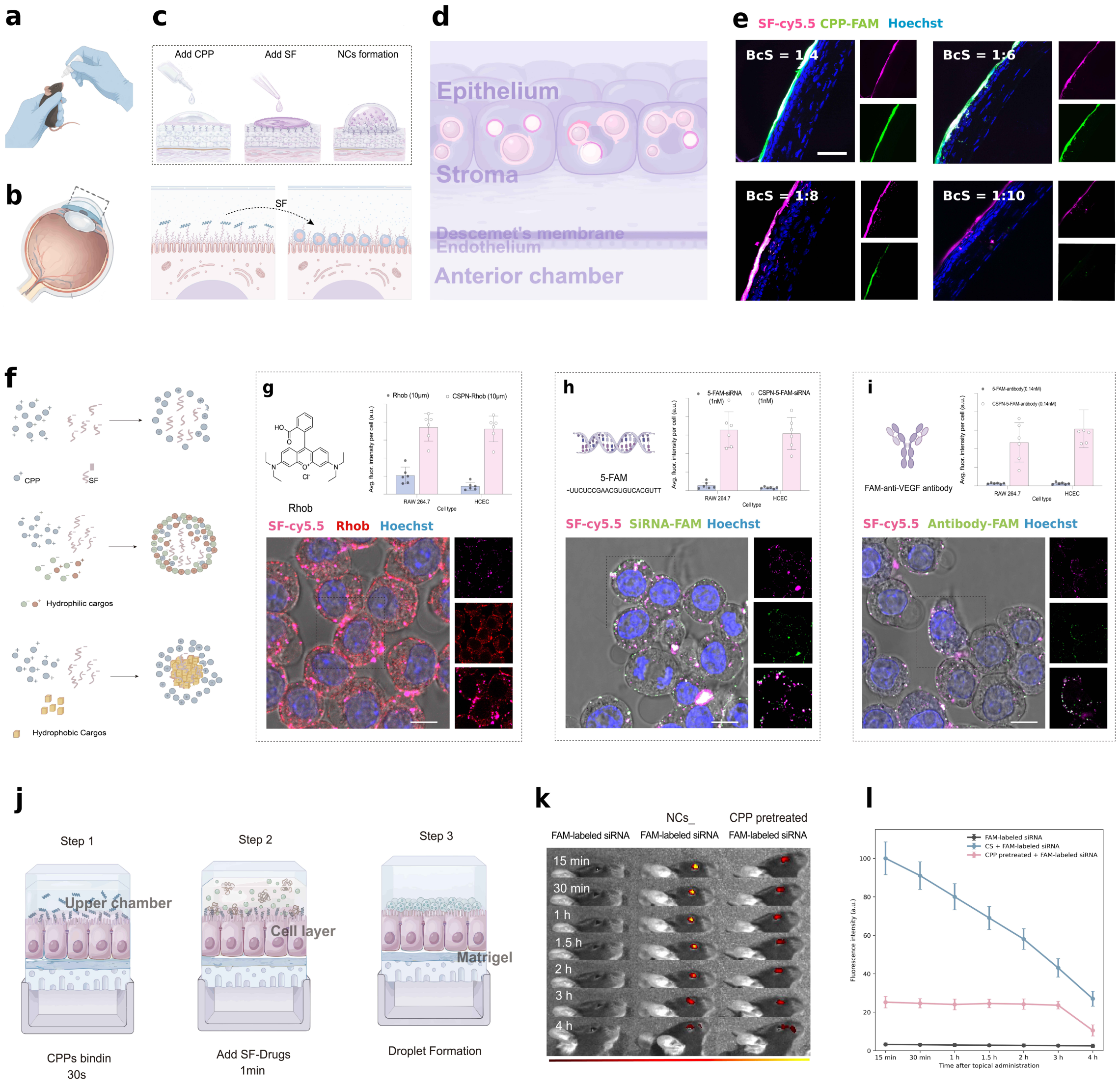


**Extended Data Fig. 6 | Ocular-surface nanocondensate assembly and cargo delivery.**

a-d, Topical administration, target region, sequential assembly and corneal transport. e, SF-Cy5.5 and CPP-FAM at the indicated BcS values. f, Cargo-loading model. g-i, Delivery of rhodamine B, FAM-siRNA and FAM-labelled anti-VEGF antibody. j, HCEC Transwell workflow. k,l, In vivo FAM-siRNA fluorescence. In e, n = 4 corneas from four mice per BcS. For g-i, n = 4 independent experiments with 30 cells per condition averaged within each experiment; two-sided one-way ANOVA used Holm-Šidák correction across formulations. For k and l, n = 6 mice per group; a two-sided mixed-effects model gave treatment-by-time $P < 0.0001$ with Holm-Šidák comparisons. Data are mean ± s.e.m. Scale bars, 10 μm (g-i) and 60 μm (e).

| Cargo category | Specific cargo | Approx. MW | Physicochemical features | EE | DL |
|---|---|---|---|---|---|
| Hydrophilic small molecule | Sulfo-Cy5.5 | ~1128 Da | Highly hydrophilic, anionic, fluorescent tracer | 38.6 | 2.4 |
| Hydrophilic small molecule | Fluorescein sodium | 376 Da | Hydrophilic, anionic | 34.2 | 2.1 |
| Hydrophilic small molecule | Rhodamine B | 479 Da | Moderately hydrophilic, aromatic structure | 46.8 | 3.5 |
| Small-molecule drug | Betaxolol | 307 Da | Moderately hydrophobic, weakly basic | 68.4 | 7.8 |
| Small-molecule drug | Dexamethasone | 392 Da | Hydrophobic, steroid structure | 73.1 | 8.9 |
| Small-molecule drug | Latanoprost | 432 Da | Highly hydrophobic, lipophilic prostaglandin analogue | 78.5 | 10.2 |
| Small-molecule drug | Timolol maleate | 432 Da | Moderately hydrophilic, ionizable | 51.7 | 5.2 |
| Small-molecule drug | Brimonidine tartrate | 442 Da | Relatively hydrophilic, cationic tendency | 43.9 | 3.8 |
| Small-molecule drug | Pilocarpine | 208 Da | Small, hydrophilic, weakly basic | 31.5 | 2.0 |
| Mechanosensitive modulator | Yoda1 | 356 Da | Hydrophobic, aromatic-rich structure | 71.6 | 8.4 |
| Nucleic acid | Negative control siRNA | ~13.3 kDa | Strongly anionic, double-stranded RNA | 88.7 | 11.8 |
| Nucleic acid | Therapeutic siRNA | ~13.5 kDa | Strongly anionic, double-stranded RNA | 86.9 | 11.3 |
| Fluorescent nucleic acid | FAM-siRNA | ~14 kDa | Strongly anionic, fluorescently labeled RNA | 84.5 | 10.7 |
| Nucleic acid | Cas9 sgRNA | ~31 kDa | Anionic, medium-sized RNA | 81.4 | 10.1 |
| Protein | BSA | 66 kDa | Water-soluble, weakly anionic protein | 58.2 | 5.9 |
| Antibody fragment | Ranibizumab Fab | ~48 kDa | Hydrophilic antibody fragment | 69.3 | 8.1 |

**Extended Data Table 1 | CPP-SF Loading Performance Classified by Cargo Hydrophilicity**
